# Readiness for behavioral change of discretionary salt intake among women in Tehran, Iran

**DOI:** 10.1101/336628

**Authors:** Nahid Kholdi, Hamed Pouraram, Ashraf Pirasteh, Mitra Abtahi

## Abstract

**Background:** It is vitally important to take into consideration women’s role in dietary pattern choice and family food management. Since women’s readiness for dietary behavioral change can be one of the most effective fundamental measures for preventing chronic diseases in developing countries, the present study is aimed to determine the readiness for behavioral change in voluntary salt intake as well as its determinants among women living in Tehran.

**Materials and methods:** The present cross-sectional study was conducted on 561 women referring to the women care units across city of Tehran. In this regard, demographic information of the participants was collected. The self-administered questionnaire included assessment of nutrition-related knowledge on salt intake and its association with diseases, discretionary salt intake, stages of change, and self-efficacy of women. In addition, the logistic regression test was used to determine the predictors of women’s readiness for behavioral change in voluntary salt intake.

**Results:** 40% women had someone in the family who had such a limitation (salt intake-limited exposure group), while 81.6% always or often added salt to their foods. Moreover, one-third of the participants were in the stage of pre-contemplation and 41.2% were in the stage of preparation for reducing salt intake. Stage of change increased with an increase in the self-efficacy score (r=0.42, p<0.001). Self-efficacy and salt intake-limited exposure were the two most important determinants of the women’s readiness for behavioral change in voluntary salt intake, respectively: (OR=1.1 95% CI: 1.06-1.14 p<0.001; OR=1.58, 95% CI: 1.03-2.42 p<0.038)

**Conclusions:** Results of the present study showed that increased self-efficacy is associated with higher levels of behavioral change among women. Since self-efficacy is very important for initiating and maintaining the behavioral change, women’s empowerment for reducing salt intake necessitates putting the emphasis on increased self-efficacy as well as community-based nutritional interventions.

## Introduction

Behavior change is a process which occurs in individuals with different levels of motivation and readiness for change [1–3]. Changing health-related behaviors can significantly affect some of the most important causes of death and diseases. Behavior plays a pivotal role in health. Evidence suggests that different behavioral patterns are deeply rooted in sociocultural conditions and depend on cultural background. Currently, behavior change interventions have a high potential for changing the current pattern of diseases. It is difficult to change the genetic pre disposition to diseases and social conditions, at least in the short term [4].

Studies which assess individuals’ readiness to change health behaviors reveal that over one-third of people are at the pre-contemplation stage and few (8-14%) at the preparation stage [5–7]. Moreover, increase in age, level of education, and self-efficacy; and the existence of chronic diseases such as hypertension which requires the limitation of salt intake, have been associated with placing individuals at higher levels of change [5, 7–8].

Hypertension is among the main risk factors for cardiovascular diseases, especially myocardial infarction, stroke, and congestive heart failure [9]. Based on the report by the Institute for Health Metrics and Evaluation of Iran, hypertension is the second risk factor among the 10 main risk factors of increased DALYs in 2010. According to this report, hypertension is among the most important causes of cardiovascular diseases [10]. Statistics show that cardiovascular diseases have the highest rate of mortality in Iran, similar to other countries. However, it is in the third rank of disease burden [11]. The main reason for increased prevalence of hypertension could be the daily growth in the elderly population, and increase in risk factors, such as unhealthy eating habits (increased salt intake, saturated fats, etc.), decreased physical activity, overweight, and stress. One of the most important factors in unhealthy eating habits is the salt intake [12].

Based on the most recent study (2014) on adults in Tehran, Iran, daily salt intake was 9 g for men and 6.96 g for women, with 53.6% being discretionary salt intake or salt intake in cooking or at the table [13]. At present, the per-capita salt intake in Iran is 10-12 g per day, higher than the amount recommended by the World Health Organization (WHO), i.e. less than 5 g per day. In other words, Iranians consume salt in their diet 5.2 times more than others [14]. A study reported mean salt intake to be 10.3 g per day in urban and rural areas of Ilam Province, Iran [15]. Moreover, it is 11.47 g per day for adults in Isfahan, Iran [16]. In terms of salt intake, Iran is similar to Denmark (7.1 g per day in women), China (12 g per day), Spain (9.8 g per day), and Japan (7.8 g per day) [17–20]. Contrary to most developed countries in which the majority of salt intake (75%) results from processed food [21–22], 50 to 60% of daily salt intake in Iran is the salt added in cooking or at the table [13,23]. In developing countries, most salt intake is the salt added in cooking or at the table. To reduce salt intake, behavioral interventions are much more effective and cost-effective than population-level interventions [24–25].

At present in Iran, reduction of salt intake is among the major priorities of planners, policy makers, and experts of the health services system [23]. On the other hand, small decreases in salt intake are associated with a significant reduction in blood pressure among those with hypertension or normal blood pressure [26–28]. That is why the WHO has recommended the reduction of daily intake of salt to 5 g for adults [29]. Studies show that, in most developing countries including Iran and China, approximately 75-80% of the total salt intake is the salt added in cooking and at the table [2]. Therefore, the major solution must focus on discretionary salt intake, with a change in dietary habits and promotion of nutrition literacy and attitude for selecting low-salt foods must be placed on the agenda so that the level recommended by WHO, i.e. less than 5 g per day, can be achieved [3, 30].

Evidence suggests that individuals benefits from reducing salt intake [26, 28]. Reducing salt intake is an easy, beneficial, independent, and low-cost method for reducing the burden of diseases, decreasing related costs, and maintaining health, and is the most effective preventive approach in most countries [31–34]. Therefore, national intervention programs, e.g. educational programs with the cooperation of the Ministry of Health, households, and related organizations, seem necessary for reducing discretionary salt intake. Considering these problems, it is vital to offer effective interventions. Of course, the use of interventional approaches for reducing discretionary salt intake requires the consideration of individuals’ readiness for change. No such study has yet been conducted in Iran on the reduction of salt intake. Few studies have been conducted in Iran on discretionary salt intake, and no information is available on Iranian women’s readiness for changing this behavior. Thus, it seems necessary to conduct studies in order to examine women’s readiness for changing the behavior of discretionary salt intake.

In the Iranian society, women have the most important role in nutrition planning and preparation of food for the family. They have the vital role of dietary patterns choice and shaping the taste of the family [35]. Mothers affect the dietary experience of family members, especially young children, and teach them the preference for the taste of salt [36]. They are responsible for cooking and controlling the amount of salt added to the food. Also, they have the important role of managing the food the family consumes. Thus, they were selected as representatives of the whole family.

The present study aimed to determine the readiness for changing the behavior of discretionary salt intake and its determinants in women residing in Tehran capital city in Iran.

## Materials and methods

Theoretical Basis of Research: The trans-theoretical model of stages of change shows the time- and motivation-related aspects of change. This model presupposes that everyone passes the stages of pre-contemplation, contemplation, preparation, action, and maintenance.

Pre-contemplation: In this stage, individuals do not think about changing their behavior in the foreseeable future which is usually defined as the following six months. In this stage, individuals have no intention to change their behavior, but are aware of the problem or behavior.

Contemplation: Individuals think about a change in the foreseeable future but not immediately, defined as 1-6 months. In this stage, they are aware of and think about and reexamine their behavior, but have not yet made a decision.

Preparation: Individuals plan for changing their behavior in the near future which is often defined as the next month. In this stage, they decide to change their behavior.

Action: Individuals have significantly changed their lifestyle in the past 6 months. Since action is visible, a change in behavior is often considered synonymous with action.

Maintenance: Individuals maintain change for some time, usually six months or more. In this stage, the behavior is stable and permanent and individuals try to prevent regression [1–3].

The model of stages of change have been employed in numerous diet-related studies in order to determine the stages of change in diet behavior, including the reduction of fat intake and increasing fiber, fruit, vegetable, and dairy intake, and its efficiency has been confirmed. Therefore, diet-related behavioral interventions are more effective if they are based on theories of health-related behavior change [30, 37–45].

### Study Design

The present cross-sectional (descriptive-analytic) study was conducted on 561 women visiting women’s care units in Tehran, Iran, selected through convenience sampling.

Inclusion criteria: Women who:

1. were willing to participate in the study;
2. were minimum 18 years of age;
3. were literate;
4. prepared food for the family;

Exclusion criteria:

1. Pregnant or breastfeed
2. have a low-sodium diet prescribed by doctors of dietitians;

### Data Collection Tool

In this study, a researcher-made questionnaire was used that included 4 sections in the fields of nutritional knowledge, discretional salt intake, women’s readiness to change behavior and self-efficacy. The questions in this questionnaire were based on the aims of the study, review of the studies, specific cultural and social conditions of Iranian women.

Demographic questionnaire included age, occupation, and level of education. In terms of occupation, women were divided into two groups: employed and home-makers. In terms of the level of education, considering the diversity in this factor, women were divided into three groups of below high school diploma (elementary and high school education), high school diploma, and university education in order to check the relationship between this factor and the qualitative variables of the study.

Self-administered questionnaire consisted of several section. In the first section measuring women’s nutritional knowledge was regarding the relationship between high salt intake and diseases. Each correct response received the score of 1 [46]. The other question was related to the presence of someone in the family who had limitations on salt intake. The answer “yes” showed the salt intake-limited exposure group in the family.

The second section focused on the discretionary salt intake. Over 60% of salt intake in Tehran is the salt added in cooking or at the table [13]. Also, it is difficult to precisely measure salt intake [6, 47]. Therefore, in order to prevent the recall bias and since self-reported avoidance of salt intake has a high correlation with the actual behavior [48], the habit of adding salt in cooking and at the table was questioned. Respondents could select one of the options of “always”, “often”, “sometimes”, “rarely”, or “never” (respectively scored 1 to 4) for each item on salt intake in cooking or at the table. To classify the responses, the answers given to these questions were congregated as “salt users” or “non-salt users”. Non-salt users were those who never, rarely, or sometimes added salt in cooking or at the table. Salt users where those who had selected the answers “always” or “often” [49, 50].

The third section determined the readiness for changing salt intake which was designed at the scale of stages of change based on the trans-theoretical model using the questionnaire used by Newson e al [6]. With this scale, the intention and decision of individuals regarding discretionary salt intake were determined by selecting one of the five stages of pre-contemplation, contemplation, preparation, action, and maintenance. To determine the predictors of readiness for changing the behavior of discretionary salt intake, stages of change were divided into three categories of pre-contemplation (not ready for change), ready for change (including contemplation and preparation), and action (change has occurred; including action and maintenance).

The fourth section was on women’s self-efficacy for reducing salt intake, including 6 questions scored on a five-point Likert scale from “Not at all sure” to “Completely sure” (scored 1 to 5, respectively). Minimum and maximum possible scores were 6 and 30, respectively. Questions in this section were designed based on the Persian adaptation of the General Self-Efficacy Scale (GSES) [51] and Self-Efficacy in Nutritional Behavior Scale (SENBS) [52].

In the pilot study conducted to determine the ease of understanding the self-efficacy questionnaire, a sample of 15 women completes the questionnaire and expressed their opinions regarding their understanding of items. In the final version, changes were applied which included the adaptation of phrases to participants and creating an appropriate framework for them.

To determine the reliability of the instrument and internal consistency of the self-efficacy items, 30 women completed the questionnaire and Cronbach’s alpha of 0.914 showed the internal consistency of this scale.

To determine the face and content validity of the questionnaire, a panel of experts (10 experts and professors) was used and their opinions regarding the clarity and appropriateness of questions for the objectives were applied.

In the data collection procedure, first explanations were given by the researcher regarding study objectives, confidentiality of data, and that no names or address had to be written on the questionnaires. Then, eligible women entered the study if they were willing to participate.

## Data Analysis

Data were analyzed in SPSS 16. Descriptive data are presented in absolute and relative frequency. The Kolmogorov-Smirnov test was used to determine the normality of quantitative data. The relationship between personal-social variables, knowledge, and self-efficacy with dependent variables (discretionary salt intake and stages of change in salt intake behavior) was assessed using Spearman’s correlation and chi-squared test. Logistic regression was employed to determine the predictors of readiness for changing the behavior of discretionary salt intake. In all tests, the significant level was <0.05. This study was approved by the Committee of Ethics in Research, Shahed University (IR.Shahed.REC.1394.280).

## Results

Mean age of women was 36.21±10.1 years, ranging from 18 to 60 years. Most women (36.2%) belonged to the age group of 30 to 39 years. Older women had lower levels of education (r=0.287, p<1.001). The level of education of 46.5% of women was university, and 38.8% (218 women) had high school diploma. Also, 66.6% of women were homemakers. Employed women had levels of education 3-fold higher than that of homemakers (p<1.001). Relative frequency of discretionary salt intake in cooking and at the table based on women’s characteristics are presented in Table 1.

**Table 1.** Relative frequency of discretionary salt intake in cooking and at the table based on women’s characteristics.

|  | Salt intake in cooking |  | Salt intake at the table |  |
| --- | --- | --- | --- | --- |
|  | Yes<br>n=458 | No<br>n=103 | Yes<br>n=128 | No<br>n=433 |
| Age (years)* |  |  |  |  |
| <30 | 86.1 | 13.9 | 29.5 | 70.5 |
| 30-39 | 81.1 | 18.2 | 20.7 | 79.3 |
| ≥40 | 77.6 | 22.4 | 19.3 | 80.7 |
| level of education** |  |  |  |  |
| Below high school diploma | 74.4 | 25.6 | 34.1 | 65.9 |
| High school diploma | 81.2 | 18.8 | 20.2 | 79.8 |
| University | 84.3 | 15.7 | 21.5 | 78.5 |
| Occupation |  |  |  |  |
| Homemaker | 82.6 | 17.4 | 23 | 77 |
| Employed | 79.7 | 20.3 | 22.5 | 77.5 |
| Salt intake- limited exposure *** |  |  |  |  |
| yes | 77.6 | 22.4 | 24.7 | 75.3 |
| no | 84.3 | 15.7 | 21.6 | 78.4 |
| Knowledge level |  |  |  |  |
| Desirable | 84.2 | 15.8 | 22.3 | 77.7 |
| Undesirable | 78.5 | 21.5 | 23.5 | 76.5 |
| Self-efficacy score**** |  |  |  |  |
| **Mean± SD** | 18.94± 6.3 | 22.45± 6.3 | 15.05± 5.7 | 20.92± 5.9 |

\*Age and salt added at the table (chi-squared test, p=0.047)
\*\* Level of education and salt added at the table (chi-squared test and p=0.029)
\*\*\* Salt intake- limited exposure group and salt added in cooking (Fisher's exact test, p=0.046)
\*\*\*\* Self-efficacy score and salt intake in cooking and at the table (Mann-Whitney U test, p<0.001 for each case)

40% women had someone in the family who had such a limitation (salt intake-limited exposure group). Compared to younger women, more women aging 40 years or older belonged to the salt intake-limited exposure group (p=0.045). The knowledge of women regarding the relationship between salt intake and diseases was undesirable in 44.7% of cases (i.e. they had answered less than half of the questions correctly), and only 28% of women had a desirable knowledge of this matter. Mean score of self-efficacy for reducing discretionary salt intake was 19.58±6.4.

### Discretionary salt intake in cooking

In cooking, over half of the women (52.6%), 163 women (29.1%), and 39 women (6.9%) always, often, and never added salt to food, respectively. Adding salt to food while cooking had a significant correlation with the salt intake-limited exposure (Fisher’s exact test, p=0.046). The ratio of women who exposure salt intake limitation and added salt to food (77.6%) was less than the ratio of women who did not exposure such a limitation and added salt to food (84.3%). Mean score of self-efficacy was higher in women who did not add salt in cooking than those who did (22.4±6.3 vs. 18.9±6.2, respectively, p<0.001).

### Discretionary salt intake at the table

Here, 298 women (53.1%), 24.1%, and 8.9% (50 women) never, sometimes, and always added salt to food at the table, respectively. Level of education and age were correlated with adding salt to food at the table. Adding salt to food at the table was less in women with university education (21.5%) than those with an education level below diploma (34.1%) (p=0.029). Moreover, women aging 40 years or above added salt to food at the table less than women aging 30-39 and below 30 years (respectively 19.3 vs. 20.7 and 29.5%, p=0.047). Mean score of self-efficacy was higher in women who did not consume salt at the table than those who did (p<0.001) (Table 1). A positive and significant correlation was observed between salt intake in cooking and at the table (r=0.194, p<0.001); 69.2% of women who never added salt to food in cooking did not consume salt at the table.

### Total discretionary salt intake (in cooking & at the table)

Table 2 shows the total discretionary salt intake based on demographic and other characteristics of women. In general, 66% of women were salt users. Age and mean self-efficacy score were significantly correlated to the total salt intake (p=0.048 and p<0.001, respectively).

**Table 2.**
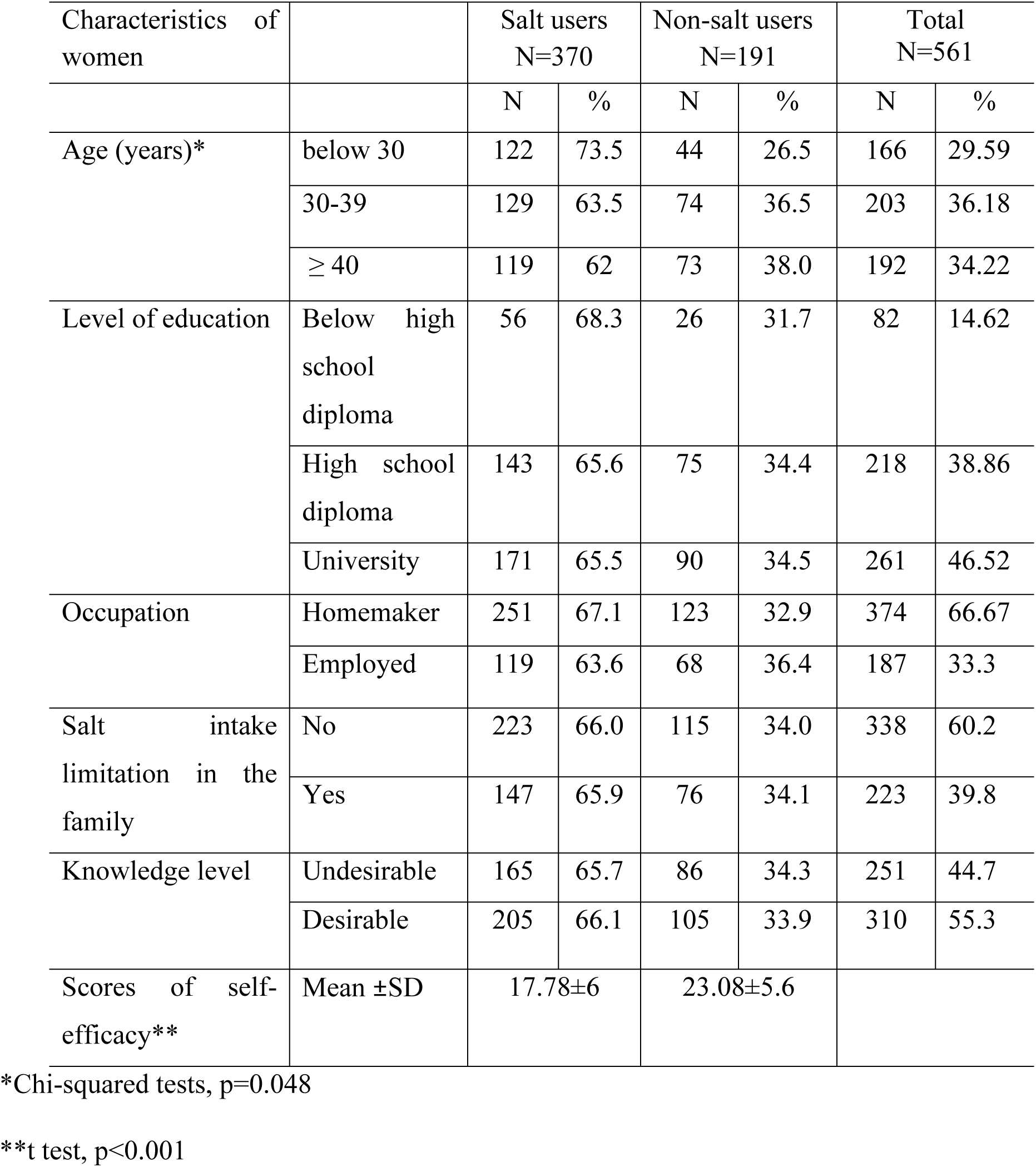
Discretionary salt intake based on the personal-demographic characteristics of women visiting women’s care units in Tehran.

### Stages of behavior change of discretionary salt intake

In terms of the behavior changing of discretionary salt intake stages, about one-third of women (31.9%) were in pre-contemplation, following by 22.3% in contemplation, 18.9% in preparation, 8.2% in action, and 18.7% in maintenance.

Table 3 presents the relationship between stages of changing the behavior of discretionary salt intake in terms of pre-contemplation, readiness (contemplation and preparation), and action (action and maintenance) with women’s characteristics. Women belonging to the age group of over 40 years were in the stage of preparation more than the two other age groups (respectively 45.3 vs. 40.4% in the age group of 30-39 years and 37.3% in the age group of <30 years) (p<0.02).

**Table 3.**
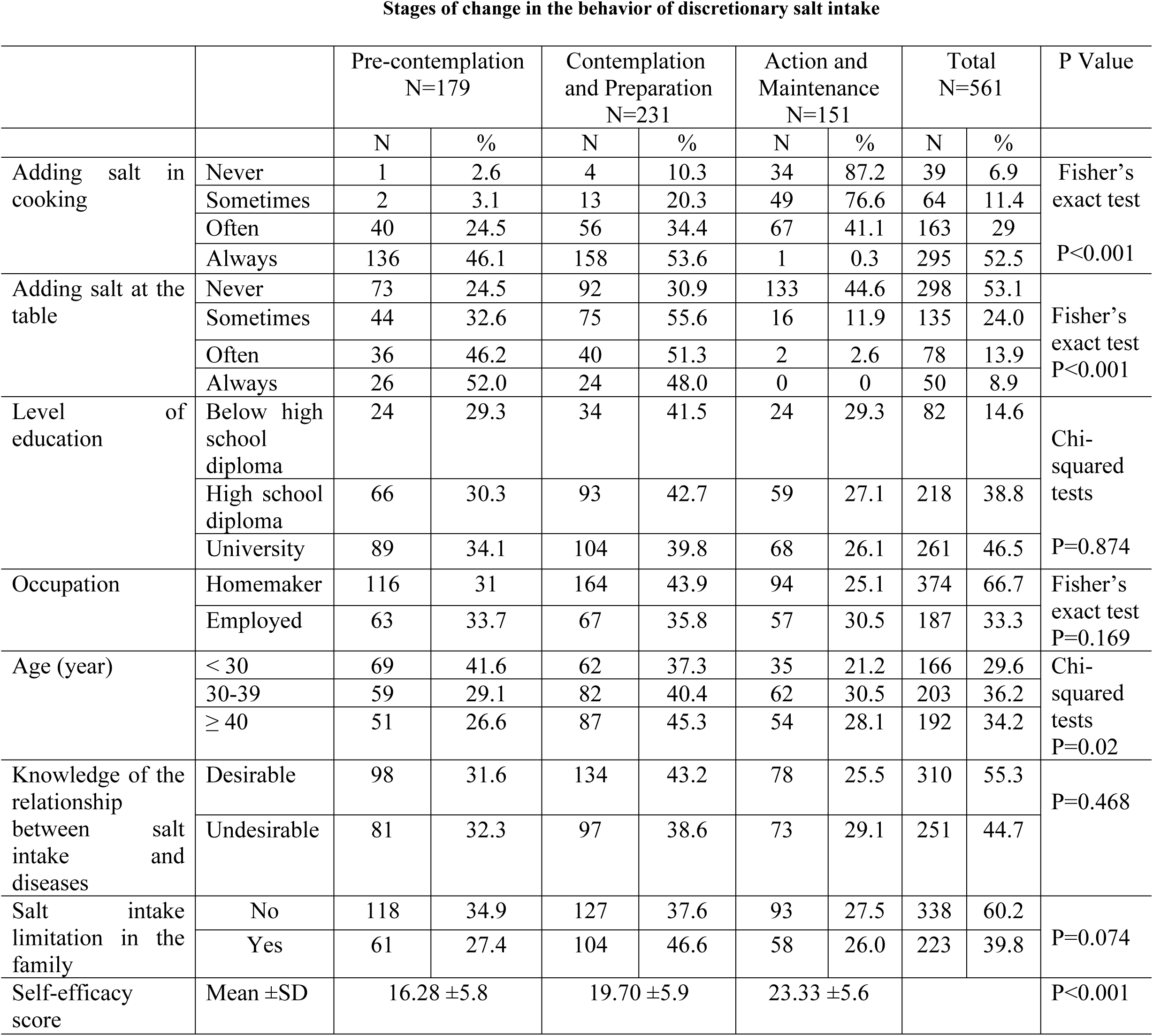
Frequency distribution of stages of changing the behavior of discretionary salt intake in women visiting women’s care units in Tehran based on studied variables.

**Stages of change in the behavior of discretionary salt intake**
|  |  | Pre-contemplation<br>N=179 |  | Contemplation<br>and Preparation<br>N=231 |  | Action and<br>Maintenance<br>N=151 |  | Total<br>N=561 |  | P Value |
| --- | --- | --- | --- | --- | --- | --- | --- | --- | --- | --- |
|  |  | N | % | N | % | N | % | N | % |  |
| Adding salt in<br>cooking | Never | 1 | 2.6 | 4 | 10.3 | 34 | 87.2 | 39 | 6.9 | Fisher's<br>exact test<br>P<0.001 |
|  | Sometimes | 2 | 3.1 | 13 | 20.3 | 49 | 76.6 | 64 | 11.4 |  |
|  | Often | 40 | 24.5 | 56 | 34.4 | 67 | 41.1 | 163 | 29 |  |
|  | Always | 136 | 46.1 | 158 | 53.6 | 1 | 0.3 | 295 | 52.5 |  |
| Adding salt at the<br>table | Never | 73 | 24.5 | 92 | 30.9 | 133 | 44.6 | 298 | 53.1 | Fisher's<br>exact test<br>P<0.001 |
|  | Sometimes | 44 | 32.6 | 75 | 55.6 | 16 | 11.9 | 135 | 24.0 |  |
|  | Often | 36 | 46.2 | 40 | 51.3 | 2 | 2.6 | 78 | 13.9 |  |
|  | Always | 26 | 52.0 | 24 | 48.0 | 0 | 0 | 50 | 8.9 |  |
| Level of<br>education | Below high<br>school<br>diploma | 24 | 29.3 | 34 | 41.5 | 24 | 29.3 | 82 | 14.6 | Chi-squared<br>tests<br>P=0.874 |
|  | High school<br>diploma | 66 | 30.3 | 93 | 42.7 | 59 | 27.1 | 218 | 38.8 |  |
|  | University | 89 | 34.1 | 104 | 39.8 | 68 | 26.1 | 261 | 46.5 |  |
| Occupation | Homemaker | 116 | 31 | 164 | 43.9 | 94 | 25.1 | 374 | 66.7 | Fisher's<br>exact test<br>P=0.169 |
|  | Employed | 63 | 33.7 | 67 | 35.8 | 57 | 30.5 | 187 | 33.3 |  |
| Age (year) | < 30 | 69 | 41.6 | 62 | 37.3 | 35 | 21.2 | 166 | 29.6 | Chi-squared<br>tests<br>P=0.02 |
|  | 30-39 | 59 | 29.1 | 82 | 40.4 | 62 | 30.5 | 203 | 36.2 |  |
|  | ≥ 40 | 51 | 26.6 | 87 | 45.3 | 54 | 28.1 | 192 | 34.2 |  |
| Knowledge of the<br>relationship<br>between salt<br>intake and<br>diseases | Desirable | 98 | 31.6 | 134 | 43.2 | 78 | 25.5 | 310 | 55.3 | P=0.468 |
|  | Undesirable | 81 | 32.3 | 97 | 38.6 | 73 | 29.1 | 251 | 44.7 |  |
| Salt intake<br>limitation in the<br>family | No | 118 | 34.9 | 127 | 37.6 | 93 | 27.5 | 338 | 60.2 | P=0.074 |
|  | Yes | 61 | 27.4 | 104 | 46.6 | 58 | 26.0 | 223 | 39.8 |  |
| Self-efficacy<br>score | Mean ±SD | 16.28 ±5.8 |  | 19.70 ±5.9 |  | 23.33 ±5.6 |  |  |  | P<0.001 |

No significant relationship was observed between the level of education, occupation, knowledge regarding the relationship between salt and diseases, and salt intake-limited exposure in the family on the one hand, and stages of change on the other. However, a significant relationship existed between stages of change and adding salt in cooking (p<0.001) and at the table (p<0.001). Women who did not add salt in cooking or at the table were at higher stages of change (respectively 56.2 and 99% in maintenance). The stage of change moved forward with increasing the score of self-efficacy (r=0.42, p<0.001). A positive correlation was observed between self-efficacy score and categorized levels of change (r=0.42 and p<0.001).

To determine the factors associated with women’s readiness for behavior change, predictor variables, including age, level of education, occupation, salt intake-limited exposure in the family, knowledge level regarding the relationship between salt intake and diseases, and self-efficacy score were entered into logistic regressions. Results showed that self-efficacy and salt intake-limited exposure are the most important factors determining women’s readiness for changing the behavior of discretionary salt intake compared to women in the pre-contemplation stage (OR=1.1 95% CI: 1.06-1.14 p<0.001; OR=1.58, 95% CI: 1.03-2.42 p<0.038) respectively.

## Discussion

The present study determined readiness for changing the behavior of discretionary salt intake in adult women residing in Tehran and its determinants, including age, occupation, level of education, knowledge level regarding the relationship between salt intake and diseases, self-efficacy score, and salt intake-limited exposure in the family as the most important determinants of women’s readiness for changing the behavior of discretionary salt intake. Results revealed that the majority of women were at the pre-contemplation stage. A limited number of participants were in the maintenance stage and had a significantly higher self-efficacy score compared to those in the pre-contemplation stage.

In the present study, women over 40 years of age consumed less salt at the table and had a more advanced level of readiness compared to the other age groups. Age is a demographic characteristic which determines discretionary salt intake. Results can be interpreted by assuming that, as age increases, individuals focus more on their health and healthy eating habits.

Level of education is an indicator of the socioeconomic status and helps a healthy diet. In the present study, the level of education had a significant correlation with the amount of salt added at the table, and women with university education consumed less salt at the table than women with an education level below high school diploma. Chen Ji [53] showed that those with a lower level of education consume more salt. Similarly, Jeong showed that those with low income and education level consume more salt [54]. It is clear that a low level of education is a serious limitation for achieving dietary knowledge and selecting healthy dietary behaviors [55]. The role of women is more important considering the pivotal role in buying the ingredients, preparing food, regulating the diet, and dietary choices [56].

In the present study, occupation as another indicator of socioeconomic status showed no significant correlation with levels of changing the behavior of discretionary salt intake. Chen Ji [53] showed that the level of salt intake was higher in those with lower levels of occupation.

These factors include a range, from individual determinants to environmental, social, and cultural characteristics. They direct individuals to attach importance to the type and quality of food based on economic conditions of the family, ensuring health, and meeting the needs.

A significant positive correlation was observed between stages of change and self-efficacy scores. In other words, self-efficacy scores increased as individuals advanced in the stages of change towards action and maintenance. In the action and maintenance stage, 87% of participants never added salt in cooking. Moreover, 45% of women in this stage never added salt to food at the table.

The study by Newson [6] showed that 22% of the sample often or always added salt to food before tasting it and 58% never added salt to food at the table. Over one-third (34%) of participants were in pre-contemplation and 28% in maintenance stages, and only 8% of the sample were at the stage of readiness for changing the behavior of discretionary salt intake, inconsistent with the results of this research.

Vander Veen [5] used the stages of change model to show that 32% of participants were at the pre-contemplation, 14% at the dynamic (contemplation, preparation, and action), and over half of them (54%) at maintenance stages. Discretionary salt intake in cooking or at the table was lower in the maintenance group than dynamic and pre-contemplation groups.

Results of a study by Ni showed that only 38% of patients who were aware of reducing salt intake actually practice it. Knowing that salt intake must be limited but not acting upon it shows that these individuals are at preparation, contemplation, or pre-contemplation stages of change [57]. Another important point is that, despite having knowledge regarding the importance and benefits of a healthy dietary pattern, practices differs [5, 6, 57]. This shows that specific measures compatible with the social context must be taken by policymakers to change this behavior among women, and education alone does not suffice in planning. Measures such as correction of the dietary environment at home may help.

A quasi-experimental study in Malaysia used the stages of change model to evaluate the behaviors of patients with hypertension in relation to doing regular exercises, reducing salt intake, and increasing the consumption of fruits and vegetables. Results revealed that patients in action and maintenance groups had significantly reduced salt intake compared to those in pre-contemplation and preparation groups.

In recent study by Jeong, the most important aspect of readiness for changing a behavior was a high self-efficacy score [54]. Participants considered environmental support and motivation as the most important factors leading to behavioral change, consistent with the results of Chen [53].

Assessing Iranian women’s readiness for change is the first step towards the assessment and promotion of food and nutrition literacy. Readiness for changing behaviors is a novel concept which receives considerable attention today. However, few studies have been conducted on this issue among women.

According to studies, the assessment of readiness for changing behaviors has the best results when the context is defined well. In the present study, women’s readiness to change the behavior was assessed using a valid questionnaire which included different dimensions of behavior change at home.

## Limitations

This questionnaire needs further corrections before being used for extensive society-level studies. A strong validation process and a larger sample size are required.

## Conclusion

The evaluation of readiness for changing a behavior is a novel concept in society-level dietary interventions. These instruments and questionnaires can be easily implemented by experts and health workers. Therefore, it can be used in large society-level studies after corrections and final approval.

Results of the present study showed that the evaluation of the stage of readiness for changing dietary behaviors can be a useful tool in interventional studies which aim to change dietary behaviors since it provides the opportunity to identify readiness for change in different social groups.

## Acknowledgments

The present study could never be brought into practice without cooperation of the participating women. Besides, the authors would like to appreciate the authorities of the women care units across Tehran who provided the ground for conducting the present research project.

## Author Contributions

**Conceptualization:** KhN PaH PA AM.

**Formal analysis:** KhN PaH PA AM.

**Methodology:** PA AM KhN PH.

**Supervision:** KhN PaH PA.

**Writing - original draft:** AM PaH.

**Writing - review & editing:** KhN PA.

